# A model of autonomous interactions between hippocampus and neocortex driving sleep-dependent memory consolidation

**DOI:** 10.1101/2022.01.31.478475

**Authors:** Dhairyya Singh, Kenneth A. Norman, Anna C. Schapiro

## Abstract

How do we build up our knowledge of the world over time? Many theories of memory formation and consolidation have posited that the hippocampus stores new information, then “teaches” this information to neocortex over time, especially during sleep. But it is unclear, mechanistically, how this actually works — how are these systems able to interact during periods with virtually no environmental input to accomplish useful learning and shifts in representation? We provide a framework for thinking about this question, with neural network model simulations serving as demonstrations. The model contains hippocampus and neocortical areas, which replay memories and interact with one another completely autonomously during simulated sleep. Oscillations are leveraged to support error-driven learning that leads to useful changes in memory representation and behavior. The model has a non-Rapid Eye Movement (NREM) sleep stage, where dynamics between hippocampus and neocortex are tightly coupled, with hippocampus helping neocortex to reinstate high-fidelity versions of new attractors, and a REM sleep stage, where neocortex is able to more freely explore existing attractors. We find that alternating between NREM and REM sleep stages, which alternately focuses the model’s replay on recent and remote information, facilitates graceful continual learning. We thus provide an account of how the hippocampus and neocortex can interact without any external input during sleep to drive useful new cortical learning and to protect old knowledge as new information is integrated.

## Introduction

Building our knowledge of the world over time requires the ability to quickly encode new information as we encounter it, store that information in a form that will serve us well in the long term, and carefully integrate the new information into our existing knowledge structures. These are difficult tasks, replete with pitfalls and trade-offs, but the brain seems to accomplish them gracefully. The Complementary Learning Systems (CLS) framework proposed that the brain achieves these feats through a division of labor across two interacting systems: The hippocampus encodes new information using a sparse, pattern-separated code, supporting rapid acquisition of arbitrary information without interference with existing neocortical knowledge (1). The hippocampus then gradually “teaches” its newly encoded experiences to the neocortex, which uses overlapping, distributed representations adept at representing the structure across these experiences, resulting in the construction of semantic knowledge over time.

CLS considered the possibility that sleep may be a useful time for this teaching to occur, and other perspectives have also focused in on this idea, given the strong coupled dynamics and parallel replay that occurs in these areas during sleep (2–4). Extant theories of consolidation have focused particularly on Stages 2 and 3 of non-Rapid Eye Movement (NREM) sleep, when nested oscillations associated with memory replay — hippocampal sharp wave ripples, thalamocortical spindles, and neocortical slow oscillations — drive especially strong hippocampal-cortical interaction (2–11). These dynamics appear to be causally involved in memory consolidation: Manipulations that enhance hippocampal-cortical synchrony during sleep benefit memory (12–16).

Not all theories agree on whether offline hippocampal-cortical interactions serve to increase the relative reliance on neocortex for episodic memories (17, 18), but most theories agree that the hippocampus helps to build and shape semantic representations in neocortex (19, 20), and these theories often assign a central role to active processing during sleep (21, cf. 22). The core ideas, shared across several perspectives, that we adopt here are: 1) During sleep, the hippocampus actively helps to build neocortical semantic representations of information it has recently encoded, and 2) this process involves a transformation of memories, not simply a transfer of memories from hippocampus to cortex. We think these processes can usefully be referred to as “systems consolidation” (23), even though the way they are conceptualized here deviates in several respects from Standard Systems Consolidation Theory (17).

Despite all of this evidence and theorizing that the hippocampus “teaches” the neocortex new information in order to build up semantic representations during sleep, it is unclear mechanistically how this might actually occur. How can brain regions interact autonomously, with no input from the environment, to produce useful learning and reshaping of representations? How does the brain move from one memory to another offline, and which of these offline states does it learn from, and how? It does not seem likely that simple strengthening of existing connections through Hebbian learning will be sufficient — a more sophisticated restructuring of representations seems to be at work. The original formulation of the CLS framework (1) did not address these questions, since there was no true implemented hippocampus; as in many models of replay, the order and nature of inputs to cortex were engineered by hand. Here, we aim to fill this gap with a model that shows how an implemented cortex and hippocampus can interact, replay, and learn autonomously.

We will address two additional related limitations of the original CLS framework, namely that its solution to continual learning was too slow, and that it relied on the implausible assumption of a stationary environment (24). Hippocampal replay of an episode — for instance, a first encounter with a penguin — was hypothesized to be interleaved over days, weeks, and months with a stationary distribution of bird input from the environment, allowing careful integration of the lone penguin encounter with the general structure of birds. This strategy is slow because it relies on these continued reminders over time about the distribution of information from the environment. But sleep-dependent memory consolidation, including integration of new information with existing knowledge, can occur quite quickly, over one night or several nights of sleep (6, 7, 25–28). The CLS framework fails in non-stationary environments because the environment no longer provides those required reminders of old information. Humans do not have this problem: We can, for example, speak one language much of our lives and then move to a new place where we encounter only a new language, without forgetting the first language. Norman, Newman, & Perotte (2005) proposed that alternating NREM and REM sleep stages across the course of a night may be the key to solving these problems (24). The hippocampus and neocortex are less coupled during REM (2), potentially providing an opportunity for neocortex to visit and explore its existing attractor space. This could serve to provide the reminders about remote knowledge needed to avoid new information overwriting the old, without the slow wait for the environment to provide those reminders.

We will present simulations that demonstrate 1) how the hippocampus can begin to autonomously teach neocortex new information during NREM sleep, and 2) how alternating NREM/REM sleep stages allows for rapid integration of that new information into existing knowledge. Our model architecture includes our previously-developed model of the hippocampus, C-HORSE (Complementary Hippocampal Operations for Representing Statistics and Episodes; 27, 28), as well as a neocortical layer as the target of consolidation (Fig 1A). Our learning scheme leverages oscillations during sleep to support error-driven learning (24). The model’s units employ a rate code, which means that oscillations do not have a true frequency that would correspond to particular frequencies of activity in the sleeping brain. But we imagine that our NREM oscillations correspond in function to sleep spindles, when replay and enhanced plasticity have been shown to occur (31, 32), and our REM oscillations to theta oscillations, which have been associated with enhanced plasticity in that stage (24, 33, 34). We simulate the difference between NREM and REM sleep as a difference in the degree of communication between hippocampus and neocortex (2). There are of course many other important differences between NREM and REM sleep (3), but for these simulations, we aimed to isolate the contribution of this factor. Together, the simulations demonstrate how sleep can build our neocortical knowledge over time, with the hippocampus initially helping to construct neocortical representations during NREM sleep, and the hippocampus and cortex working together across NREM/REM stage alternation to integrate this new information with existing knowledge.

**Fig 1.**
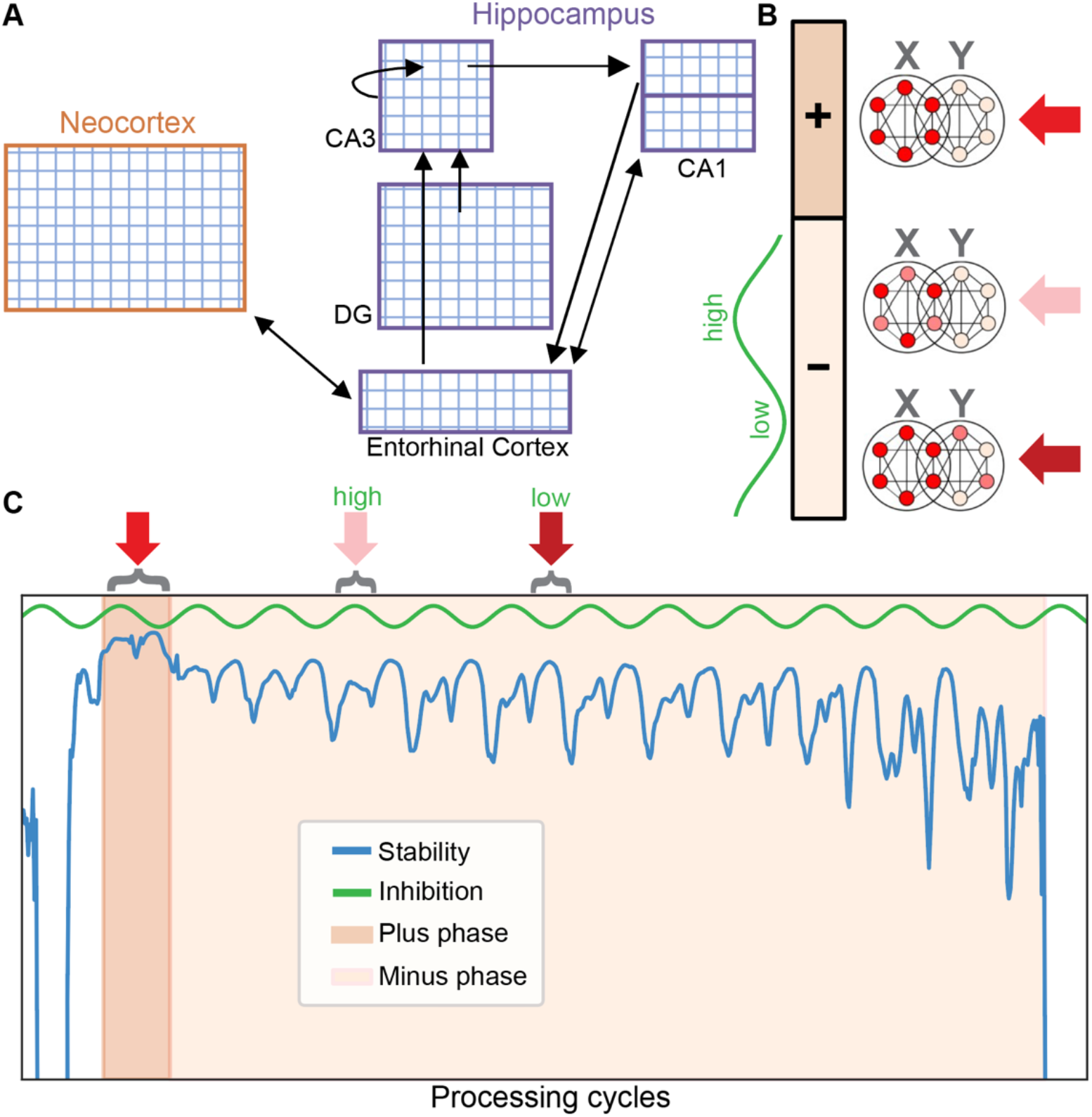
Model architecture and sleep algorithm. **(A)** The architecture included C-HORSE (purple outlines) — our model of the hippocampus, and a neocortical layer as the target of consolidation (orange outline). **(B)** When replaying attractor X, all units participating in the attractor have strong, stable activity during the plus phase. During the minus phase, phases of higher inhibition lead to lower activity, with the weakest units in attractor X dropping out, and phases of lower inhibition lead to higher activity, causing activity spreading to a nearby attractor, Y. (**C**) Stability trace for a real learning event, with background colors indicating the plus and minus phases. When the model first falls into an attractor, its activity is highly stable and triggers the plus phase. Short-term synaptic depression gradually destabilizes the attractor. The stability drop causes the plus phase to end and the minus phase begins. As the attractor further destabilizes, the minus phase ends.

## Model simulations

In the model, sleep is initiated with a single injection of noise to all units, after which all external input is silenced. The model then autonomously moves from attractor to attractor, corresponding to items learned during wake. We use stability in the model’s dynamics as a trigger to initiate a learning trial. When the model initially falls into each attractor state, activity tends to be highly stable from one processing cycle to the next, and this high stability initiates a “plus” phase (Fig 1B,1C). Short-term synaptic depression destabilizes the attractor by temporarily weakening synapses in proportion to the coactivity of their connected units. As stability drops, the model transitions into a “minus” phase. Oscillations are always present in the sleeping model, but they become more prominent in the minus phase, as synaptic depression begins to distort an attractor. These oscillations in inhibition levels reveal aspects of the attractor in need of change: when inhibition is high, the weakest units in the attractor drop out, revealing the parts of the memory that need strengthening, and when inhibition is low, nearby potentially-interfering memories or spurious associations become active, revealing competitors in need of weakening (Fig 1B, 1C).

As the attractor activity further destabilizes, the minus phase ends and the model traverses to the next attractor. Weights are updated at the end of every minus phase via Contrastive Hebbian Learning, which locally adjusts minus phase unit coactivities (product of sender and receiver activity over time) towards their plus phase coactivities (35). This serves to both strengthen weak parts of an attractor (the contrast between the plus phase and the higher inhibition moments in the minus phase) and weaken competitors (the contrast between the plus phase and the lower inhibition moments of the minus phase). This algorithm is closely related to the Oscillating Learning Rule (OLR; 24), which used oscillations to similarly distort attractors and improve memory, but the OLR requires the network to track where it is in an oscillation and change the sign of the learning rule accordingly. The current learning scheme does not require tracking this information (but does require a measure of stability).

### Simulation 1: Building neocortical representations of novel information

In this simulation, we explored how the hippocampus helps to rapidly shape neocortical representations of novel information. As a demonstration, we simulated a paradigm in which we have found sleep effects that influence behavior quickly (across a night of sleep or even a nap) and that have hallmarks of the construction of new neocortical knowledge (27, 36). In the experiment, participants were asked to learn the features of fifteen satellites belonging to three different classes (Fig 2A; 27). Each satellite exemplar had features shared with other members of the class, as well as features unique to the exemplar. As in the human participant sleep group, the model was first trained to a learning criterion of 66% and then allowed to sleep, with tests before and after sleep to assess changes in performance (Fig 2C, 3A). In this simulation, sleep corresponds to the dynamics of NREM, with tightly coupled hippocampal and neocortical areas.

**Fig 2.**
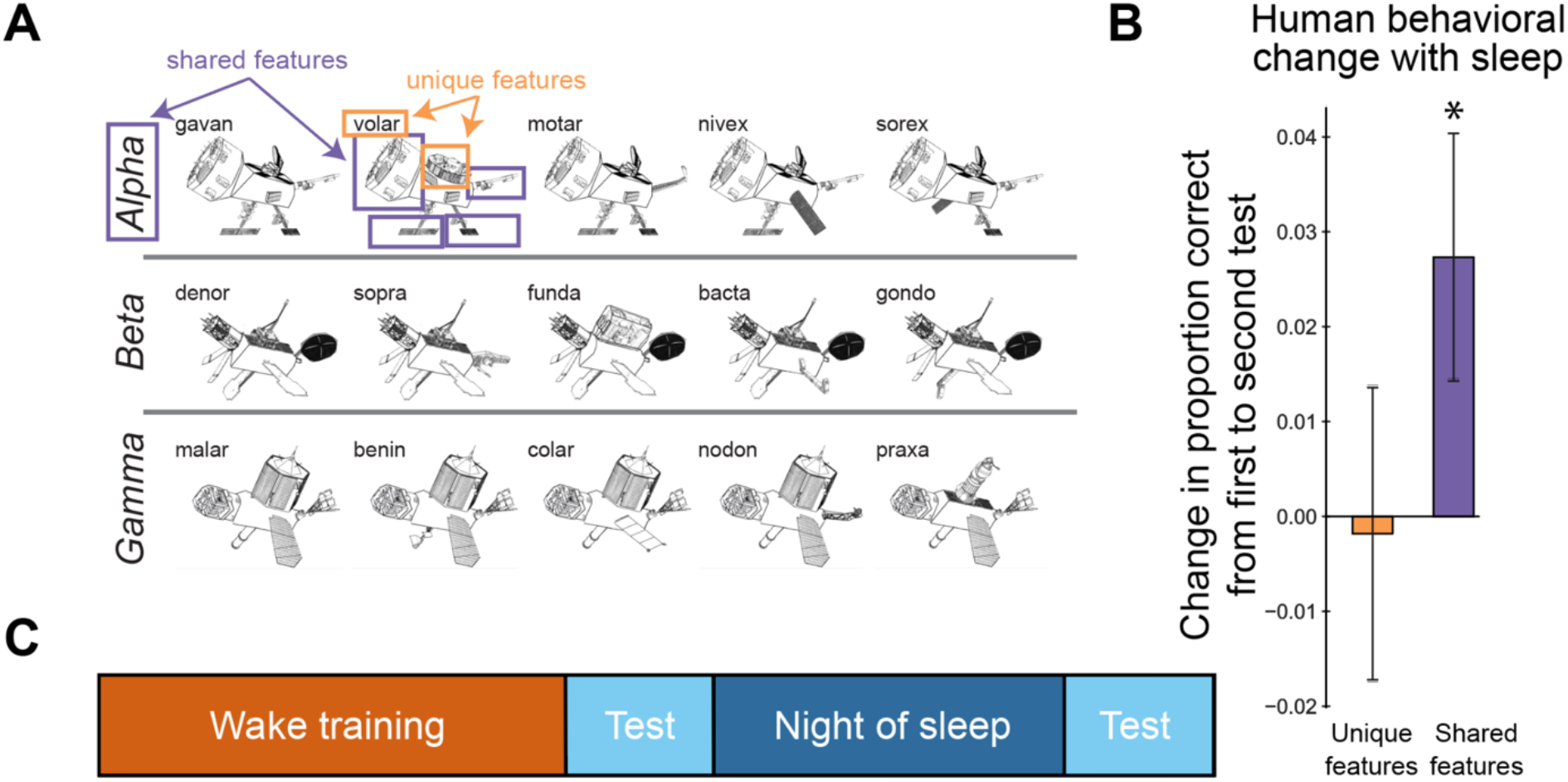
Schapiro et al. (2017) category learning paradigm and results. **(A)** Satellite exemplars studied from three categories. **(B)** Performance change for unique and shared features from pre-to post-sleep tests. *p < .05. (Results are collapsed across the frequency manipulation included in that study.) **(C)** Experimental protocol: Participants studied the satellites in the evening, had an immediate test, and were tested again 12 hours later, after a night of sleep.

**Fig 3.**
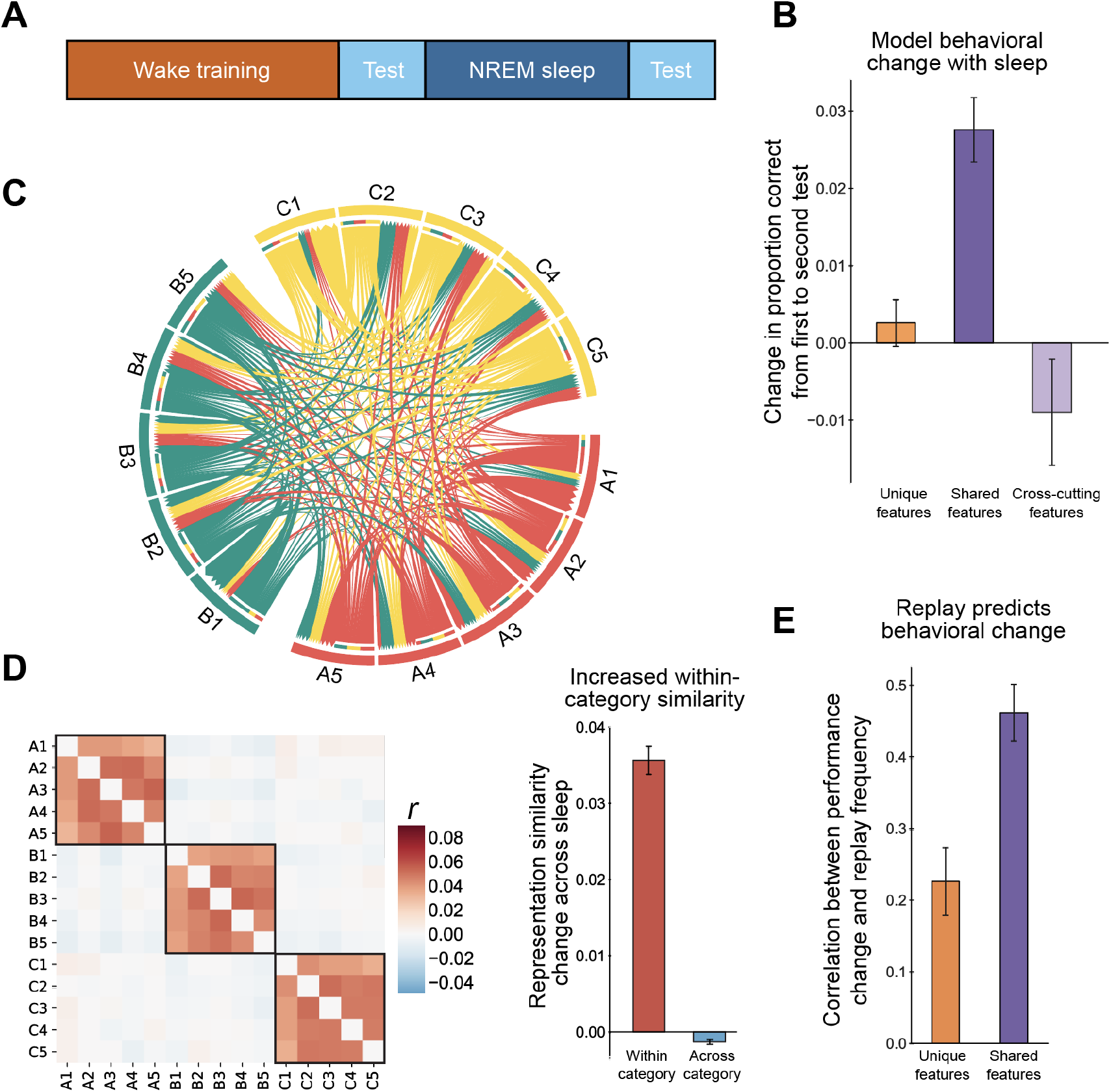
Simulation 1: Category learning consolidation across one night of sleep. **(A)** Simulation protocol. **(B)** Change in model performance from before to after sleep for unique features, shared features, and cross-cutting features. **(C)** Chord plot of replay transitions, with A, B & C indicating categories. Arrow base indicates transition initiation and head indicates termination. Arrow width indicates proportion of replays for a given satellite. The plot shows that replay was interleaved uniformly across exemplars (with a small preference for within-category transitions). **(D)** Left: Change in pairwise correlation across items from before to after sleep. Black boxes indicate within-category changes. Right: Averaged pairwise correlation change within and across categories. **(E)** Fisher-transformed correlation between exemplar replay frequency during sleep and performance change, averaged across model initializations. All error bars represent +/− one SEM across network initializations.

The model was trained during wake using feature inference, which is analogous to how humans were trained in the experiment. Each satellite’s seven features (five visual, class name, and codename) were presented on separate input/output layers (representing entorhinal cortex) as inputs to the hidden layers of the model. Each training trial consisted of presenting the model with a satellite with one feature missing, at which point the model guessed the identity of the feature. The correct answer was then clamped onto the corresponding layer. Contrastive Hebbian Learning was used to update the weights based on the difference between the model’s prediction and the correct answer (35). To better match model performance on shared and unique features during wake training, as with human participants, unique features were queried much more often. The learning rate was much higher in the hippocampus than cortex, so — while learning happened during wake across the model — behavior was initially supported almost entirely by the hippocampus. Cortical representations were on average less sparse than hippocampal representations and reflected category structure more strongly. For this paradigm, which requires integrating high level verbal and visual information, the neocortical module might correspond to an area of the brain representing consolidated semantic knowledge, like the anterior temporal lobe (37). Anterior cingulate cortex / medial prefrontal cortex is also often involved as an important neocortical target of systems consolidation (38, 39).

After the wake learning criterion was achieved, we allowed the model to go into NREM sleep, enabling hippocampal-cortical replay. The cortical learning rate was increased, inspired by evidence for enhanced plasticity in sleep (40). To isolate the impact of replay on neocortical learning, we paused hippocampal learning during sleep. The model traversed from attractor to attractor corresponding to the studied satellite exemplars. Replay occurred concurrently in the hippocampus and neocortical modules. We found that replay was interleaved across exemplars with a small preference for within-category transitions (39.2% within-category transitions relative to chance 33.3%; p < 0.001 by binomial test [*N* = 2162, *K* = 847]; Fig. 3C). We assessed performance changes as a result of NREM sleep and found that performance on shared features improved significantly (*p* < 0.001) while performance on unique features was preserved across sleep (Fig. 3B), which corresponds to the pattern observed in the human experiment shown in Fig. 2B as well as the behavioral data from our imaging study using this paradigm (36).

The preferential benefit for shared features could potentially be related to their greater frequency of occurrence, given that they appear many times across exemplars, whereas unique features only appear with one exemplar. To rule out the possibility that sleep is simply benefitting higher-frequency features, we ran a separate set of simulations in which we added an extra feature to each exemplar that appeared across categories but was matched in frequency to the shared features. We found that these “cross-cutting” features did not benefit from sleep replay (Fig 3B), suggesting that neocortical learning was sensitive to the true category structure, rather than simply frequency of feature appearance.

We assessed whether, for each simulation, the number of times a particular satellite was replayed was associated with improvement in memory for that satellite. This resulted in a correlation across 15 exemplars for each simulation, which we Fisher transformed and averaged across simulations. We found that, across simulations, there was a positive correlation between performance changes and replay frequency (Fig. 3E), which was especially strong for shared features, indicating that replay during sleep was driving performance changes. This is consistent with the results from our imaging study of this paradigm, where we found that awake replay of the satellites in the hippocampus (which we took to be representative of what would continue to happen during sleep) predicted memory improvement across sleep (36).

We next examined change in neocortical representations across sleep. We conducted a representation similarity analysis by calculating pairwise correlations of patterns of neocortical unit activity for all satellite exemplars pre- and post-sleep. We found that, from before to after sleep, there was increased within-category similarity and a small decrease in across-category similarity (Fig 3D).

The results from Simulation 1 provide a way of thinking about how the hippocampus can help the neocortex build up new semantic representations. With a fast learning rate in the hippocampus and slow rate in neocortex during wake learning, the hippocampus will be responsible for most behavior prior to sleep. Offline, the hippocampus can help the neocortex to reinstate stable versions of the new memories. Because the neocortex has more highly distributed representations than the hippocampus, it tends to find the shared structure across exemplars, resulting in improved understanding of the category structure in this paradigm. These dynamics occur completely autonomously, with three key ingredients: 1) Short-term synaptic depression causes the hippocampus and neocortex to move in tandem from one attractor to the next, resulting in interleaved replay of recent experience; 2) as synaptic depression destabilizes an attractor, oscillations increasingly distort the attractor to reveal aspects of the memory in need of change; and 3) the model learns toward highly stable states and away from the subsequent, relatively-unstable states, resulting in memory shifts and strengthening.

### Simulation 2: Integration of recent and remote knowledge

We next turned to simulating the potential role of sleep in allowing graceful continual learning. We defined two related environments, Env 1 and Env 2. Each environment contained 10 items, with seven units each. Two out of seven units overlapped between the first item in Env 1 and the first item in Env 2, between the second item in Env 1 and the second item in Env 2, and so on. There was no overlap across items within an environment. This is a variant of the classic AB-AC interference paradigm (41–44). To simulate new learning in Env 2 after having fully consolidated Env 1 in neocortex, we trained the neocortical hidden layer fully on Env 1. Next, the full network was trained on the related Env 2, with hippocampus primarily supporting performance, as in Simulation 1, given its faster learning rate. We then allowed the model to go into either an alternating sleep protocol, alternating five epochs of NREM and REM sleep, or into five sequential NREM epochs (Fig 4A). NREM was implemented as above, with strong coupled dynamics between hippocampus and neocortex, whereas REM allowed neocortex to explore its attractor space without influence from the hippocampus.

**Fig 4.**
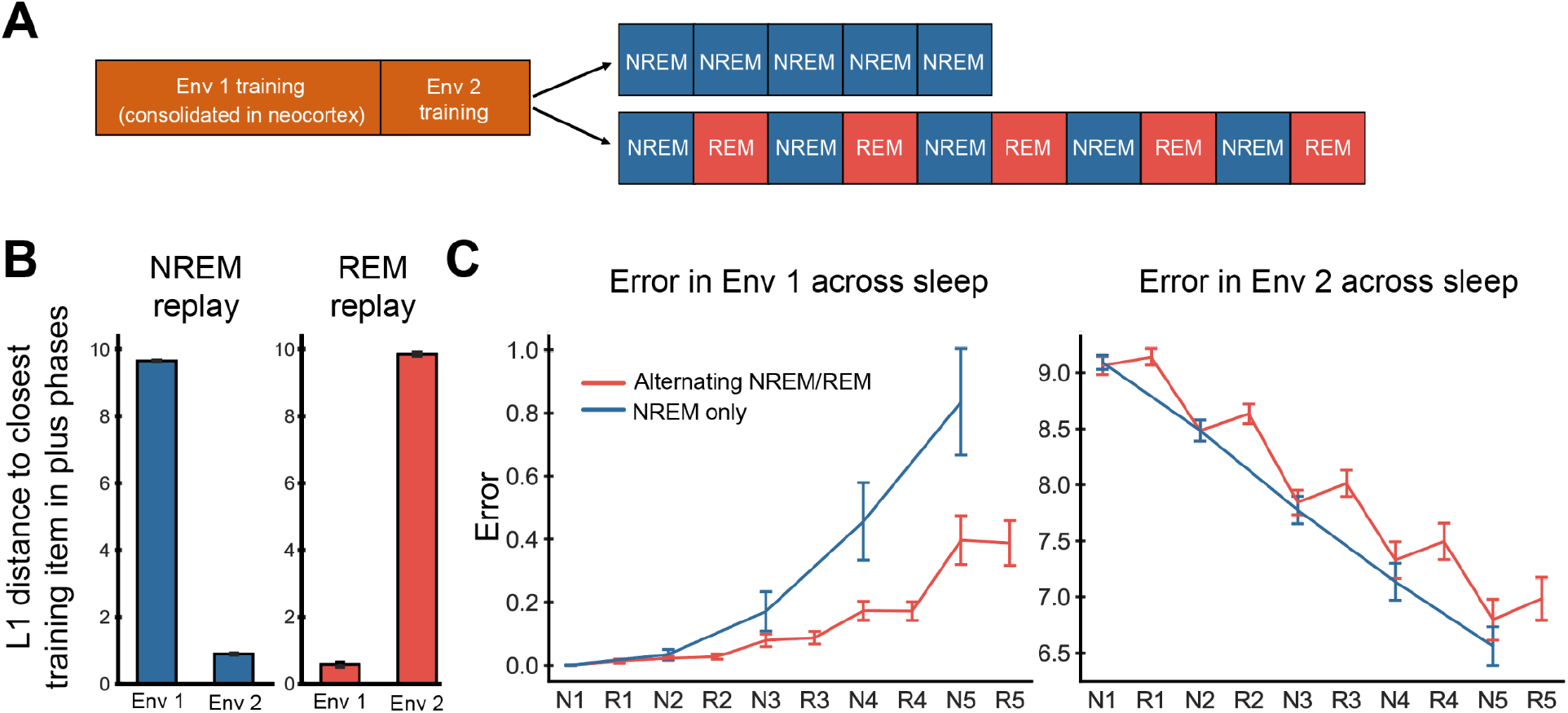
Simulation 2: Continual learning via alternating sleep stages. **(A)** Simulation protocol. **(B)** L1 distance of plus phase attractors to Env 1 & 2 item patterns in REM and NREM. Smaller distance indicates more replay of those patterns. **(C)** Error across sleep for Env 1 & 2 items for alternating NREM/REM (N/R) vs. NREM-only conditions.

We expected that Env 2 replay would unfold during NREM sleep in a manner analogous to Simulation 1, with hippocampus helping the neocortex to replay and learn the Env 2 patterns. During REM, with less influence from the hippocampus, we expected that neocortex would be able to replay the well-consolidated remote Env 1 memories. This could serve to provide reminders about Env 1 to prevent Env 2 from overwriting Env 1. We observed, as expected, that NREM tended to focus on Env 2 items (lower distance from plus phase patterns to Env 2 items; Fig 4B) whereas REM tended to focus on the remote Env 1 items.

We found that alternating NREM and REM allowed the neocortex to reduce error on Env 2 without a catastrophic increase in error on Env 1 (Fig 4C). Consecutive epochs of NREM sleep also resulted in reduction of error on Env 2 but led to a much more substantial increase in error on Env 1.

Alternating NREM and REM may thus support graceful continual learning, with NREM helping to build neocortical representations of recent information while REM acts to protect the old information from this new potential interference (45). This alternation should be especially important to the degree that new and old information overlap and therefore threaten interference; when new information is unrelated to prior knowledge (as in Simulation 1), reminders of old knowledge are likely not needed (46).

## Discussion

Our simulations demonstrate how the hippocampus can autonomously teach the neocortex new information during NREM sleep, and how alternating between NREM and REM sleep over the course of the night can promote graceful integration of new information into existing cortical knowledge. The hippocampus is implemented as our C-HORSE model, which is able to quickly learn new categories and statistics in the environment, in addition to individual episodes (29, 30, 47). We extend the model with a neocortical area that serves as the target of consolidation and with a sleep environment that allows learning and dynamics between these areas to unfold autonomously offline.

Simulation 1 demonstrates how the hippocampus can begin to shape neocortical knowledge during a first bout of NREM sleep. We think of this as the very beginning of systems consolidation, not the complete process. Still, one night of sleep (or even a nap) is sufficient to appreciably change behavior in many paradigms, including the task simulated here (27, 36). Sleep begins by seeding the network with random activity. The network then falls into an attractor that happens to resemble that activity, and moves from attractor to attractor due to a short-term synaptic depression mechanism which weakens synapses as a function of their recent activity. We find that this mechanism results in interleaved replay of attractors, which sets the stage for useful structure learning (1). When the network first falls into an attractor, activity is highly stable, and the network uses these stable states as targets to learn towards. Oscillations then distort these attractors, revealing contrasting states to learn away from — both weak parts of an attractor in need of strengthening and competing attractors in need of weakening. During NREM, the hippocampus helps the neocortex to reinstate strong versions of attractors (neocortex’s own attractors for new information would not be strong enough to support offline learning on their own), and this autonomous error-driven learning allows useful weight updates from these states. Almost all prior models of replay, including the original CLS model, do not have this fully autonomous dynamic (48, cf. 49). Typically, the model architect exerts some control over the input to the model on a given replay trial, and indeed defines what a “trial” is. However, a large part of the computational puzzle of understanding learning during sleep is understanding how the system can generate its own learning trials, and our account provides a demonstration of how this can work.

The mechanisms thus act as hypotheses for how sleep carries out autonomous replay: once the system has visited an attractor, there should be an endogenous mechanism of leaving this attractor in order to transition to the next, which we hypothesize manifests as a form of adaptation at the synaptic level. Unit level adaptation or homeostatic mechanisms may also be possible (49), though they should result in less ability to transition between closely related memories (which are likely to share units). To define offline learning trials, we hypothesize that the brain contrasts periods of stable with periods of perturbed activity, consistent with several accounts of biologically plausible error-driven learning (50–52). We further hypothesize that oscillations (ripples/spindles in NREM and theta in REM) enhance learning during sleep by distorting attractors during the unstable (minus phase) periods in ways that are informative about flaws in the network’s representation. This implies that dampening or eliminating these oscillations would reduce the efficacy of learning during sleep, consistent with findings from both NREM and REM sleep (33, 53).

In Simulation 1, as in our human experiment results, we found that these autonomous learning dynamics resulted in enhanced memory for features of exemplars shared across a category while preserving memory for exemplar-unique features. This was not due to shared features appearing more frequently in the stimulus set, as frequency-matched features that appeared across categories did not show this improvement. Sleep resulted in increased similarity for neocortical representations of items in the same category, consistent with the literature suggesting that sleep helps to extract the structure or central tendencies of a domain (39, 54–56). To the extent that detailed information is present in the hippocampus, it should be able to help neocortex learn this detailed information, and there is extensive evidence that sleep indeed benefits retention of detailed information (26, 57, 58). However, the overlapping nature of representations in the neocortex (especially relative to the hippocampus) implies that neocortex will tend to emphasize shared structure over the course of systems consolidation.

Simulation 2 proposes a role for alternating NREM and REM sleep stages in enabling successful continual learning. We simulated learning of a new environment, reliant on the hippocampus, after the neocortex had already consolidated an old, related environment. We found that the hippocampus helps to build neocortical representations of the new environment during NREM sleep (as in Simulation 1) and the neocortex then reinstates the old environment during REM sleep, allowing careful integration of the new information into neocortex without overwriting the old. Our proposal is strongly consistent with studies suggesting that memory reactivation during SWS sets up subsequent integration into existing knowledge during REM (59, 60), and more generally with the idea that both NREM and REM sleep are critical for memory integration, with the ordering of the stages being consequential (61, 62).

The simulation demonstrates how a night of sleep can serve to protect against interference even in non-stationary environments. The original CLS framework did not provide an account of this learning scenario, as it required the environment to continue to provide reminders of old knowledge, the role taken over by REM here. The sleep model of Norman et al. (24) showed how REM can provide the necessary reminder and repair function (63), but implemented continued exposure from the environment over time to learn the new knowledge, the role taken over by NREM here. Deep neural networks performing both experience replay and generative replay have had impressive success in tackling the problem of continual learning in complex nonstationary environments (64–68). Our approach is related in benefitting from offline interleaved replay, but fleshes out the autonomous interactions between the hippocampus and cortex that may support this learning, and explains how this can happen *quickly* upon transition to a new environment through the alternation of NREM and REM sleep.

Overall, we view the process of sleep-dependent memory consolidation not as a simple strengthening of individual memories or weakening of noise (69), but instead as a restructuring that acts to update our internal models of the world to better reflect the environment over time. According to our framework, new information learned over the course of one waking period will be quickly encoded by the hippocampus. We predict that the hippocampus and neocortex will then concurrently replay this information in interleaved order during NREM sleep, with the hippocampus helping the neocortex to reinstate higher-fidelity versions of recent information than the neocortex could support without hippocampal influence. This interleaved replay should support new learning in neocortex that serves to especially emphasize the structure of the new domain, increasing representational overlap between related entities. This prediction has not yet been directly tested, but it is consistent with a finding of increased representational overlap over time for related memories in medial prefrontal cortex (39). The hippocampus could help the neocortex to reinstate higher-fidelity representations of recent experience even if the hippocampus was not required for the initial learning, consistent with recent findings in humans and rodents (70, 71), as long as the hippocampus was encoding the details of the experience in parallel with other regions.

Our account also makes predictions about sleep stage contributions to memory consolidation. First, NREM sleep should more frequently visit attractors corresponding to recent experience, and REM sleep should more frequently visit attractors corresponding to remote, consolidated knowledge, though this is not absolute. This prediction is consistent with the finding that NREM dreams incorporate more recent episodic information in comparison to REM dreams, which incorporate more existing semantic information (72). There have been very few empirical demonstrations of replay during REM (73, 74), and our account suggests that this may be partly because REM is not as focused on recent experience, which is the subject of most experiments. Our account predicts that integration of new information without disruption to existing knowledge requires NREM-REM alternations. If REM is suppressed, or in particular if replay during REM is suppressed, the account predicts eventual overwriting of old information that is related to new information (Fig 2C).

Our model’s simulations are intended as demonstrations as opposed to a full account of offline consolidation, but there are exciting possible future directions, as well as complementary modeling frameworks that already help to provide a more complete understanding. For example, our current model does not simulate changes in memory across periods of wake. Often sleep studies find a marked reduction in performance with wake, including in our satellite category learning study (27). Simple reductions in performance can be simulated through interference caused by waking experience and/or decay in weight strengths over time. However, the story is not so simple, because awake replay can be beneficial for memory (75–78). It may be that awake replay works against other causes of memory degradation, and/or that awake replay does not result in the kind of lasting improvements to performance that sleep replay does; perhaps awake replay leads to short term, local improvements, but the specialized coordinated hippocampal-cortical replay dynamics that exist during sleep are needed for persistent systems-level change (79).

We have used the word “replay” in the manner it is used in the modeling literature — offline reactivation of any information previously experienced in the environment. Sometimes the word is reserved in the empirical literature for the *sequential* reactivation of states experienced in a particular order. Most of the rodent literature has studied this sequential reactivation, and it will be important for future versions of our model to simulate these kinds of sequential paradigms, taking inspiration from other modeling frameworks that have focused on simulating sequential reactivation (49, 80, 81).

Another limitation of our framework is that our use of a rate code (unit activation values that correspond to a rate of firing across time) make the modeling of oscillations more abstract, and limit the ability to explain or make predictions about detailed oscillatory dynamics across the brain. However, a biologically detailed thalamocortical neural network model has been developed to engage with these dynamics and provides important insight into how they may contribute to memory consolidation (49, 82, 83). Having these models at different levels of abstraction is useful in creating the full bridge from individual neurons to dynamics across systems to complex behavioral tasks.

Another important missing piece in the current modeling framework, which has been tackled in other frameworks (46, 48, 80, 84, 85), is an explanation of how memories are *prioritized* for replay, for example along dimensions of emotion, reward, or future-relevance (54). We and others have also found evidence for prioritization of weaker memories for offline processing (27, 36, 43, 63, 86). Prioritization of weaker information is not something that falls naturally out of the current framework; an additional tagging mechanism is needed to mark memories with higher uncertainty or error during or after learning for later replay.

Our current simulations focused on hippocampal influence on neocortical representations during sleep, but future simulations should also consider sleep-dependent learning locally within the hippocampus (34), including differential learning in the monosynaptic (MSP) and trisynaptic pathways (TSP). Some of the behavioral change seen in the sleep consolidation literature could arise from this local hippocampal learning (22).

The hippocampus module in our framework is a version of our C-HORSE model (29, 30, 47), which rapidly learns both new statistical and episodic information, in its MSP and TSP, respectively. The original CLS framework proposed that the hippocampus handles the rapid synaptic changes involved in encoding new episodes, in order to prevent the interference that would occur from attempting to make these synaptic changes directly in neocortex. The neocortex then slowly extracts the statistics across these episodes to build up semantic information over time (weeks, months, years). But we can learn new semantic and statistical information much more quickly than this (within minutes or hours) and the hippocampus is likely responsible for much of this learning (29, 47, 71). Our view is thus that the hippocampus acts as a temporary buffer for both statistics and episodes — for any novel information requiring large synaptic changes that might cause interference if implemented directly in neocortex.

The hippocampus in the original CLS account (and also in experience replay models) functions somewhat like the TSP in our hippocampus model, replaying individual episodes and exemplars with high fidelity. The MSP, however, is more sensitive to the structure across experiences (29, 47). This suggests the possibility of generalized replay, which has indeed been observed (87, 88), and which may serve to catalyze the consolidation of structured information (89).

We have modeled the hippocampus and one generic neocortical module, but of course the brain is much more heterogeneous and hierarchical than this. We could consider the TSP and the neocortex module as two points on a spectrum, with the TSP learning very quickly using orthogonalized representations, and the neocortex learning slowly using overlapping representations. The MSP has intermediate learning speed and overlap in its representations, and perhaps hierarchically organized neocortical regions form a gradient (90), with slower learning as a region gets closer to the sensory periphery — from hippocampus, to MTL cortex, to high level sensory regions, to low level sensory regions. Each region could help to train the adjacent slower region through concurrent offline replay. The hippocampus — both the MSP and TSP — may be especially important for tracking information across the time period of one day (or across one waking period, for animals sleeping in shorter bouts). Then, with the first bout of sleep, the hippocampus starts to help adjacent neocortical areas build representations of the new information from the prior waking period.

While some learning does occur in neocortex during initial waking exposure, both in our model and in the brain, it may not typically stabilize or be strong enough to support behavior prior to hippocampal influence during sleep (91–93). When information is strongly consistent with prior knowledge, there may be the possibility for small tweaks that allow direct integration into neocortical areas without the need for extensive offline restructuring (94, 95). Our account, as well as the original CLS account, predicts that with *enough* exposure, any information, even completely novel information, can eventually be learned in neocortex without hippocampal influence (perhaps explaining the high functioning of developmental amnesics; 96). But novel information can be learned much faster if it goes through the process of quick encoding in the hippocampus followed by neocortical integration offline.

In summary: The model provides a framework for understanding how the hippocampus can help shape new representations in neocortex, and how alternating NREM/REM sleep cycles across the night continues to allow additions to this knowledge as we encounter new information over time. We hope it will inspire empirical tests and provide a foundation for exploring the mechanisms underlying a range of sleep-dependent memory findings.

## Methods

We implemented our simulations in the Emergent neural network framework (97). The model consists of units with rate code activations (using the NXX1 activation function) organized into layers and connected by learnable weights. Feedforward feedback (FFFB) inhibition is implemented to simulate inhibitory network dynamics.

### Model architecture

The model architecture for both simulations consisted of the four C-HORSE hidden layers representing the DG, CA3, pCA1, and dCA1 subfields of the hippocampus, and one neocortical hidden layer (see Supplementary Material for parameter details). Previously C-HORSE has been employed to simulate hippocampal contributions to statistical learning, associative inference, and category learning (29, 30, 47) and comes from a lineage of hippocampal models of episodic memory (16, 98). We adopted the version from Zhou et al. (30), which divides CA1 into proximal and distal components.

Input/output layers in Simulation 1 consisted of seven feature layers, each corresponding to a satellite feature’s high level visual and verbal representation in Entorhinal Cortex (EC). Simulation 2 had a separate input and output layer, corresponding to superficial and deep EC layers (29, 30).

### Wake training

We departed from the Emergent default learning in using a fully error-driven learning scheme with Contrastive Hebbian Learning (CHL; 35). CHL computes errors locally but behaves similarly to the backpropagation algorithm (52, 99). In each trial, CHL contrasts two states: a “plus” phase in which targets are provided, and a “minus” state, where the model makes a guess without a target. Given a learning rate parameter ϵ and settled sender (s) and receiver (r) activations at each synapse in the last cycle of the plus (+) and minus (−) phases, each weight update (Δw) is calculated as:

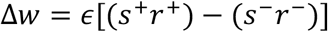

In the first simulation, the model was trained on fifteen satellite exemplars from three categories (Fig 2A). Each satellite consisted of seven features (five visual parts, a classname, and a codename) of which some were shared across exemplars from a category and others unique to each exemplar. Satellites were chosen randomly in each training trial and one feature was held out. Features were presented as one-hot inputs on the corresponding input/output feature layer and were held out at a ratio of unique:shared = 99:1 (to better match unique and shared learning speed, unique features need to be queried the vast majority of the time). Each training trial consisted of a minus and plus phase. In the minus phase the satellite was presented to the model with a feature held out and the model generated a prediction for the identity of the feature. In the plus phase, the full satellite was presented to the model. The model was trained to a criterion of 66% feature completion performance for shared and for unique features. All results are averages across 100 model initializations. All weights were randomly sampled from uniform distributions with every new initialization (means and ranges listed in Supplementary Material).

In the second simulation, each training trial minus phase involved clamping an Env 1 or Env 2 item on the input layer and requiring the model to reproduce that item on the output layer. Each environment had 10 items and each item consisted of seven units. Env 1 and Env 2 item units were chosen such that there was a 2/7 unit overlap between corresponding items from Env 1 and Env 2 (between Item 1 from Env 1 and Item 1 from Env 2, Item 2 from Env 1 and Item 2 from Env 2, and so on). There was no overlap between items from the same environment. The same set of items were used across all initializations of the model. In the plus phase, the item was clamped onto both the input and output layer. The training protocol involved first overtraining the model with only the neocortical hidden layer active on Env 1 items to simulate consolidated remote knowledge. We trained the model for thirty epochs after reaching zero error. We then switched to Env 2 training, with both the hippocampus and neocortex engaged in learning to zero error. Weight changes were calculated via CHL (Eq1) based on the coactivation differences between the plus and minus phases in both simulations. All results are averages across 30 model initializations.

### Wake testing

For both simulations, testing involved presenting the input patterns and allowing the model to generate an output using its learned weights. The model was evaluated on the distance of its output to the target. In Simulation 1, performance for the 78 shared and 27 unique features across the satellite exemplars was individually tested on feature-held-out testing trials in each testing epoch. The model’s performance on each trial was judged as correct if it generated activity on the correct units greater than a threshold of 0.5 and activity on incorrect units less than 0.5. Shared and unique proportion correct scores were then separately calculated by averaging binary correct/incorrect responses in each feature category for each model initialization. These results were then averaged across random initializations.

In Simulation 2, testing performance was calculated as the sum of squared errors in the model’s response on the output layer for each testing trial. Each testing epoch consisted of testing all ten Env 1 items and all ten Env 2 items.

### Sleep activity dynamics

Sleep epochs were initiated with a random noise injection after which all external inputs were silenced. Recirculating activity dynamics allowed the model to autonomously reinstate learned item attractors. Each synapse in the model was subject to short-term synaptic depression on its weight as a function of coactivation-induced calcium accumulation. Given *Ca*_*inc*_, *Ca*_*dec*_ time constant parameters, s sender activation, r receiver activation and wt synaptic weight, the calcium update on each processing cycle was computed as:

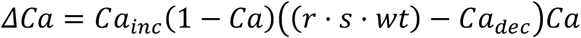

Given a calcium-based depression threshold parameter *Ca*_*thr*_and gain parameter *SD*_*gain*_, synaptic depression was computed on the weights as:

If *Ca* > *Ca*_*thr*_:

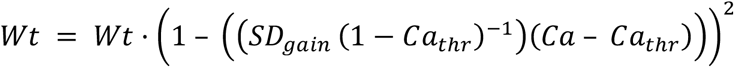

Inhibitory oscillations were implemented via a parameterized sinusoidal wave. Given the default layer FFFB inhibitory conductance Gi parameter *Gi*_*def*_, amplitude *A*, period *P*, a midline shift *S* and processing cycle *c*, the FFFB Gi parameter *Gi*_*c*_ for a given cycle for each layer was set as:

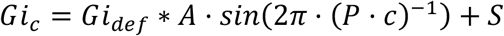

### Sleep learning

Sleep learning events were defined by the stability of model activity, calculated as the average temporal autocorrelation of layer activity in the model. Given n model layers’ activity L indexed by i, processing cycle c and the Pearson’s correlation function Corr, stability was computed as:

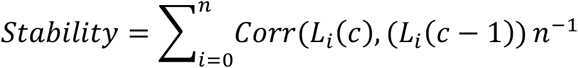

Plus phases were marked as contiguous cycles where stability was greater than a strict plus threshold (0.999965 and 0.9999 for Simulation 1 and 2, respectively). Minus phases were marked by the periods following plus phases where stability was greater than the minus threshold (0.997465 and 0.9899 for Simulation 1 and 2, respectively). Weights were updated at the end of every minus phase. Weight changes were computed using a modified version of the CHL learning rule used in wake training (Eq1). Changes were based on the difference in average minus and plus phase coactivations at each synapse:

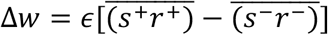

This average allows coactivity dynamics throughout the minus phase (both high and low phases of inhibition) to contribute to the contrast with the plus phase.

### Sleep stages and simulation protocols

NREM in both simulations involved all C-HORSE layers and the neocortical hidden layer being active, allowing unimpeded communication between hippocampus and neocortex. For REM sleep, C-HORSE and neocortex were disconnected (lesioned connection from entorhinal cortex to neocortex), implementing the idea that neocortex dynamics are less influenced by the hippocampus during REM. As we were focused on neocortical learning, C-HORSE projections in both sleep stages were non-learning. Only input/output layer <−> neocortical layer projections updated during sleep.

In Simulation 1, after the wake training criterion was achieved, the model executed one 30,000 cycle epoch of NREM sleep. Test epochs before and after sleep established performance change over sleep. In the second simulation, after neocortex-only Env 1 item training and full model Env 2 item training, the model either performed five alternating 10,000 cycle epochs of NREM and REM sleep, or five 10,000 cycle epochs of consecutive NREM. Test epochs established performance after every sleep epoch.

## Acknowledgments

We are grateful to James Antony, Elizabeth McDevitt, and Sharon Thompson-Schill for helpful discussions and Randall O’Reilly and Seth Herd for model implementation support. This work was supported by NIH R01 MH069456 to KAN and Charles E. Kaufman Foundation KA2020-114800 to ACS.

## Supplementary Material

Layer Parameters (Simulation 1)
| Layer | Parameter | Value |
| --- | --- | --- |
| pCA1 | # Units | 100 |
| | Inhibition | FFFB $G_i = 2.2$ |
| dCA1 | # Units | 100 |
| | Inhibition | FFFB $G_i = 2.2$ |
| DG | # Units | 225 |
| | Inhibition | FFFB $G_i = 2.8$ |
| CA3 | # Units | 144 |
| | Inhibition | FFFB $G_i = 5$ |
| Feature Layers | # Units | 6/3/90 (Visual/Classname/Codename) |
| | Inhibition | FFFB $G_i = 2.1/1.6/2.6$<br>(Visual/Classname/Codename) |
| Neocortex | # Units | 225 |
| | Inhibition | FFFB $G_i = 2.4$ |

Projection Parameters (Simulation 1)
| Projection | Parameter | Value |
| --- | --- | --- |
| Default | Weight Init Distribution | 0.25 - 0.75 |
| Default | Connectivity | Full |
| Default | Learning rate | 0.05 |
| Feature Layers -> DG | Learning rate | 0.1 |
| Feature Layers -> DG | Connectivity Sparsity | 0.6 |
| Feature Layers -> CA3 | Learning rate | 0.1 |
| Feature Layers -> CA3 | Connectivity Sparsity | 0.1 |
| DG -> CA3 | Learning rate | 0 |
| DG -> CA3 | Connectivity Sparsity | 0.1 |
| DG -> CA3 | Weight Init Distribution | 0.89 - 0.91 |
| CA3 -> CA3 | Learning rate | 0.1 |
| CA3 -> CA3 | Weight Init Distribution | 0 - 0.6 |
| Feature Layers <-> Neocortex | Learning rate | 0.0001 |

Sleep Parameters (Simulation 1)
| Projection | Parameter | Value |
| --- | --- | --- |
| Feature Layers <-> Neocortex | Learning rate | 0.03 |
| Feature Layers -> DG | Learning rate | 0 |
| Feature Layers -> CA3 | Learning rate | 0 |
| CA3 -> pCA1 | Learning rate | 0 |
| pCA1 -> Feature Layers | Learning rate | 0 |
| Feature Layers <-> dCA1 | Learning rate | 0 |
| Synaptic Depression | Inc | 0.00035 |
|  | Dec | 0.00025 |

Layer Parameters (Simulation 2)
| Layer | Parameter | Value |
| --- | --- | --- |
| pCA1 | # Units | 100 |
|  | Inhibition | FFFB Gi = 2.2 |
| dCA1 | # Units | 100 |
|  | Inhibition | FFFB Gi = 1.8 |
| DG | # Units | 225 |
|  | Inhibition | FFFB Gi = 2.8 |
| CA3 | # Units | 144 |
|  | Inhibition | FFFB Gi = 5 |
| Input/Output | # Units | 120 |
|  | Inhibition | FFFB Gi = 3.2/2.8 (Input/Output) |
| Neocortex | # Units | 144 |
|  | Inhibition | FFFB Gi = 2 |

Projection Parameters (Simulation 2)
| Projection | Parameter | Value |
| --- | --- | --- |
| Default | Weight Init Distribution | 0.25 - 0.75 |
| Default | Connectivity | Full |
| Default | Learning rate | 0.05 |
| Input -> DG | Learning rate | 0.1 |
| Input -> DG | Connectivity | 0.6 |
| Input -> CA3 | Learning rate | 0.1 |
| DG -> CA3 | Learning rate | 0 |
| DG -> CA3 | Connectivity Sparsity | 0.1 |
| DG -> CA3 | Weight Init Distribution | 0.89 - 0.91 |
| CA3 -> CA3 | Learning rate | 0.1 |
| CA3 -> CA3 | Weight Init Distribution | 0 - 0.6 |
| Input -> Neocortex | Learning rate | 0.05/0.0001 (Env1/Env2) |
| Neocortex -> Output | Learning rate | 0.05/0.0001 (Env1/Env2) |

Sleep Parameters (Simulation 2)
| Projection | Parameter | Value |
| --- | --- | --- |
| Input <-> Neocortex | Learning rate | 0.05 |
| Neocortex -> Output | Learning rate | 0.05 |
| pCA1 -> Neocortex | Learning rate | 0 |
| dCA1 -> Neocortex | Learning rate | 0 |
| Feature Layers -> DG | Learning rate | 0 |
| Feature Layers -> CA3 | Learning rate | 0 |
| CA3 -> pCA1 | Learning rate | 0 |
| pCA1 -> Output | Learning rate | 0 |
| Input <-> dCA1 | Learning rate | 0 |
| Synaptic Depression | Inc | 0.0009 |
|  | Dec | 0.0005 |

